# The Influence of Ligands on AlphaFold3 Prediction of Cryptic Pockets

**DOI:** 10.64898/2026.01.04.697564

**Authors:** Maria Lazou, Felix Tuchscherer, Sandor Vajda, Diane Joseph-McCarthy

## Abstract

Cryptic pockets are binding sites that are formed or exposed upon a conformational change. They represent an important class of potentially druggable binding sites. Reliably predicting cryptic pockets capable of binding ligands, however, remains a challenge. Herein we examine the use of AlphaFold 3 (AF3) for generating realistic conformational ensembles that include known cryptic pockets. We find that AF3 is generally able to reproduce the scale of conformational change required for cryptic site formation. When given a cryptic-site ligand for the protein, AF3 predominantly predicts conformations competent to bind the ligand in the cryptic site; without the ligand, conformations lacking the cryptic pocket generally dominate. While the results may reflect a bias toward memorized structural priors, the level of detrimental memorization appears to be limited. We also show that the choice of the ligand can significantly impact the predictions, and that AF3 is able to produce models with the ligand correctly positioned. Variability in ligand position, however, suggests that generating ensembles of co-folded predictions is critical to enhancing the likelihood of obtaining a correct binding mode. Overall, AF3-generated protein-ligand structural ensembles have potential utility in cryptic-site drug discovery, and they can reveal ligands likely to bind to those sites.

## Introduction

Cryptic sites are hidden sites that become exposed upon a relatively small conformational change from the native structure. They are important in that they can expand the range of druggable targets.^1–4^ Specifically, cryptic sites can extend an active site, act as allosteric sites, or serve as a binding site for PROTACS and/or molecular glues^5^.

As a result, considerable effort has been devoted to predicting cryptic sites through a variety of physics-based methods.^6–10^ Others have sought to augment physics-based computational approaches with artificial intelligence. ^11,12^ While significant progress has been made, challenges remain.^13^ In addition, it has long been known that proteins exist as conformational ensembles and that protein flexibility is key to understanding protein function.^14,15^ Related to this it has been shown that AlphaFold2 (AF2) models of proteins can have similar accuracy to solution NMR structures which capture a certain level protein flexibility.^16^

AlphaFold 3 (AF3), compared to AF2, has the capability of predicting ligand bound vs. ligand unbound structures.^17,18^ It operates by utilizing a single unified diffusion-based approach that treats all molecular components as part of an integrated system from the start. In this study, we examine the ability of AF3 to predict the ligand bound conformation of cryptic sites both with and without the ligand present. This comparison allows us to assess the ability of AF3 to model induced-fit and conformational selection mechanisms, and to determine whether it can reveal cryptic sites even in the absence of explicit ligand cues.

This work builds on our previous study looking at the ability of AF2 to predict cryptic pockets^19^. We and others have shown that AF has a strong bias towards the dominant conformation and has memorized alternative conformations.^20–23^ AF2 can nonetheless confidently predict different structures from the same MSA.

Herein we compare the distribution of conformations generated by AF3 with and without an input ligand to the distribution of bound and unbound structures in the protein data bank (PDB) for a test set of known cryptic pockets. We also examine the accuracy of the ligand placement by AF3, the local structure surrounding the ligand, and the effect of running AF3 with different ligands. A detailed analysis is provided.

## Results and Discussion

### Agreement between AF3 prediction and PDB distributions

For cryptic sites, that are observed upon conformational change, AF3 with multiple seeds tends to find the dominant bound conformation when predicting the protein structure with the ligand present and the dominant unbound conformation when predicting the protein structure without the ligand. The distribution of structural models produced using AF3 with a ligand is generally in agreement with the distribution of bound structures found in the protein data bank (in 16 out of 16 cases) while the distribution produced by AF3 without a ligand is generally in agreement with the distribution of unbound structures in the PDB (in 13 out of 16 cases). In 10 out of 16 cases, the predicted distributions match both the bound and unbound PDB distributions. The test set is described in Table 1 and the distributions and root mean square deviation (RMSD) plots are shown in Figure 1-3 and Figure S1. This observation is consistent with previous results showing that AF2 predictions are driven by a memory of what is in the PDB^24^. Here the distributions are sharpened somewhat by AF3 relative to what is observed in the PDB. It is clear that the same MSA is capable of predicting a range of conformations at the scale (side chain movements, loop movements, and helical shifts) necessary for cryptic site formation.

**Figure 1.**
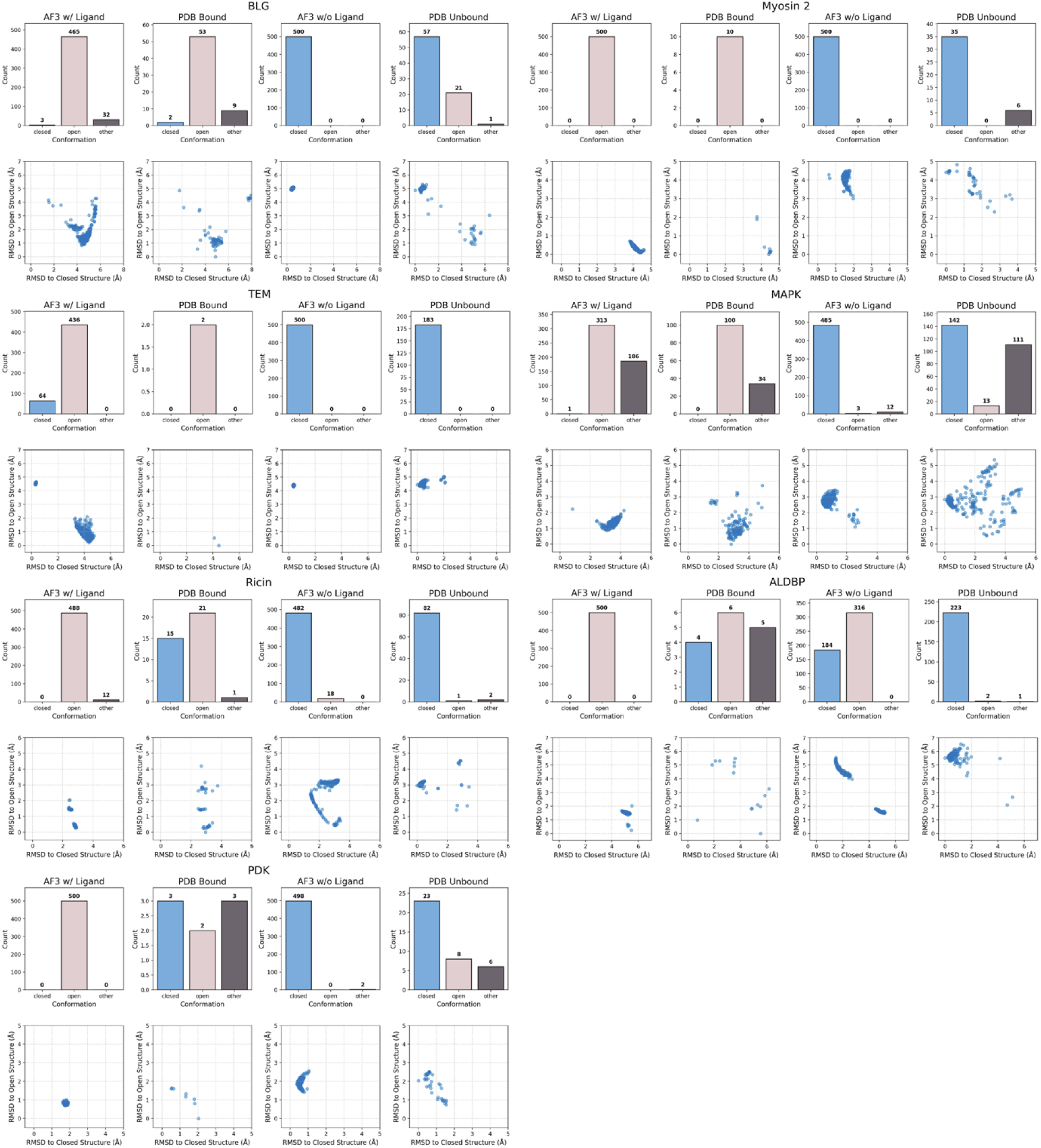
Comparison of bound and unbound structures with AF3 predictions with and without a ligand for Group 1a proteins, where the PDB distributions reveal an open and close state and AF3 accurately predicts both states. Distributions of the structures (from left to right for each protein, AF3 with ligand, PDB bound, AF3 without ligand, PBD unbound) and RMSDs of the moving residues relative to the open and closed reference structures are shown.

**Table 1.** Proteins used in the study.

| Protein | AF PDB ID | FTMove PDB ID | Closed Ref. PDB ID | Open Ref. PDB ID | Bound PDB ID | Bound Struct | Ligand ID | Moving Segment | Site Explanation |
| --- | --- | --- | --- | --- | --- | --- | --- | --- | --- |
| Bovine $\beta$ - Lactoglobulin (BLG) | 6GE7.A | 1BSQ.A | 1BSQ.A | 1GX8.A | 1GX8.A | Open | RTL | Ile84 - Asn90 | Overlaps with active site |
| Myosin II | 2AKA.A | 2AKA.A | 1D0X.A | 1YV3.A | 1YV3.A | Open | BIT | Leu262 | Fully separate cryptic |
| TEM β-lactamase (TEM) | 1PZO.A | 1PZO.A | 1JWP.A | 1PZO.A | 1PZO.A | Open | CBT | Ala217- Leu225 | Adjacent to active |
| Mitogen-Activated Protein Kinase (MAPK) | 2NPQ.A | 2ZB1.A | 2ZB1.A | 2NPQ.A | 2NPQ.A | Open | BOG | Trp197- His199 | Fully separate cryptic |
| Ricin | 1RTC.A | 1RTC.A | 1RTC.A | 1BR6.A | 1BR6.A | Open | PT1 | Tyr80 | Overlaps with active site |
| Adipocyte Lipid Droplet Binding Protein (ALDBP) | 1ALB.A | 1LIC.A | 1ALB.A | 3HK1.A | 3HK1.A | Open | B64 | Phe57 | Overlaps with active site |
| Pyruvate Dehydrogenase Kinase (PDK) | 2BU8.A | 2BU8.A | 2BU8.A | 2BU2.A | 2BU2.A | Open | TF1 | Leu29- Phe31 | Fully separate cryptic |
| Glutamate receptor 2 (GluR2) | 1MY0.B | 1MY0.B | 1MY0.B | 1N0T.D | 1N0T.D | Open | AT1 | Gly136- Ser142 | Overlaps with active site |
| Heat Shock Protein 90 (Hsp90) | 1YES.A | 1YES.A | 2QFO.B | 1YET.A | 1YET.A | Open | GDM | Gly108- Thr109 | Overlaps with active site |
| K-Ras | 4EPW.A | 4EPW.A | 4EPR.A | 4EPV.A | 4EPV.A | Open | 0QX | Met67, Tyr71 | Fully separate cryptic |
| Ribonuclease A (RNase A) | 1RHB.A | 2W5K.B | 1RHB.A | 2W5K.B | 2W5K.B | Open | NDP | His119 | Overlaps with active site |
| Androgen receptor (AR) | 2AX9.A | 2AX9.A | 2AX9.A | 2PIQ.A | 2PIQ.A | Open | RB1 | Lys720, Met734 | Fully separate cryptic |
| AmpC Beta-Lactamase (AmpC) | 2BLS.B | 2BLS.B | 2BLS.B | 3GQZ.A | 3GQZ.A | Open | GF7 | Asn289- Leu293 | Adjacent to active |
| cAMP-dependent protein kinase (PKA) | 2GFC.A | 2GFC.A | 2GFC.A | 2JDS.A | 2JDS.A | Open | L20 | Thr51 - Arg56 | Overlaps with active site |
| Thrombin | 1GHY.H | 1HAG.E | 3BEI.B | 1GHY.H | 1GHY.H | Closed | 121 | Gly216 - Cys220 | Overlaps with active site |
| β-Secretase (BACE) | 3IXJ.C | 1W50.A | 3IXJ.C | 1W50.A | 3IXJ.C | Closed | 586 | Gly66 - Glu77 | Overlaps with active site |

### Accuracy of AF3 predictions

It is typically for most cryptic sites to be formed in the open conformation. In our dataset, for 15 of the 16 proteins, the bound reference structure is in the open conformation and the unbound reference structure is in the closed conformation. For BLG, Myosin 2, TEM, MAPK, Ricin, ALDBP, and PDK, AF3 correctly predicts the bound distribution with the ligand as predominantly open and the unbound distribution without the ligand as predominantly closed (Group 1a proteins, Figure 1). As an example, the top ranked model produced for BLG with the ligand (with a ligand RMSD to X-ray pose of 1.17 Å) is shown in Figure 4. Top ranked models for the other proteins in Group 1a are shown in Figure S1.

For GluR2, Hsp90, and K-Ras, AF3 predicts the bound distribution with the ligand correctly but not the unbound distribution (Group 1b proteins, Figure 2). For GluR2, AF3 without the ligand incorrectly and almost exclusively predicts the open conformation, only two closed conformations (out of 500) are generated. However, many of the closed conformations in the PDB “unbound” distribution (210/340) actually have glutamate bound in the cryptic pocket. These are considered unbound in our test set because the glutamate does not clash with the moving segment, although it is possible that the closed position is more preferred when glutamate is bound at the site. For Hsp90, in the reference structures there is a flexible loop (residues 106 to 113 which includes the moving segment residues) that causes a break in a helix. A closer look at the distribution of structures in the PDB reveals that Hsp90 adopts at least three different conformations: one with the helix-loop-helix, another with a helix, and a third in which the protein has undergone a more significant conformational change. All three of these conformations are represented in the bound and unbound structures to some degree. AF3 with and without the ligand, models the whole segment (residue 100 to 124) as a helix with a slight kink in it. K-Ras is a structure with many very flexible regions. The helix spanning residues 65 to 74 (which includes the two residues in the moving segment) shifts from the closed to the open conformation. In the bound structures, the range of conformations of the helix is centered around the open conformation, while in the unbound structures, there is a continuous distribution from the closed to open conformations. When clashing with the reference ligand was added as a criterion for being closed (likely needed since the moving segment is in a helix), the PDB distributions for bound and unbound were open and closed as anticipated.

**Figure 2.**
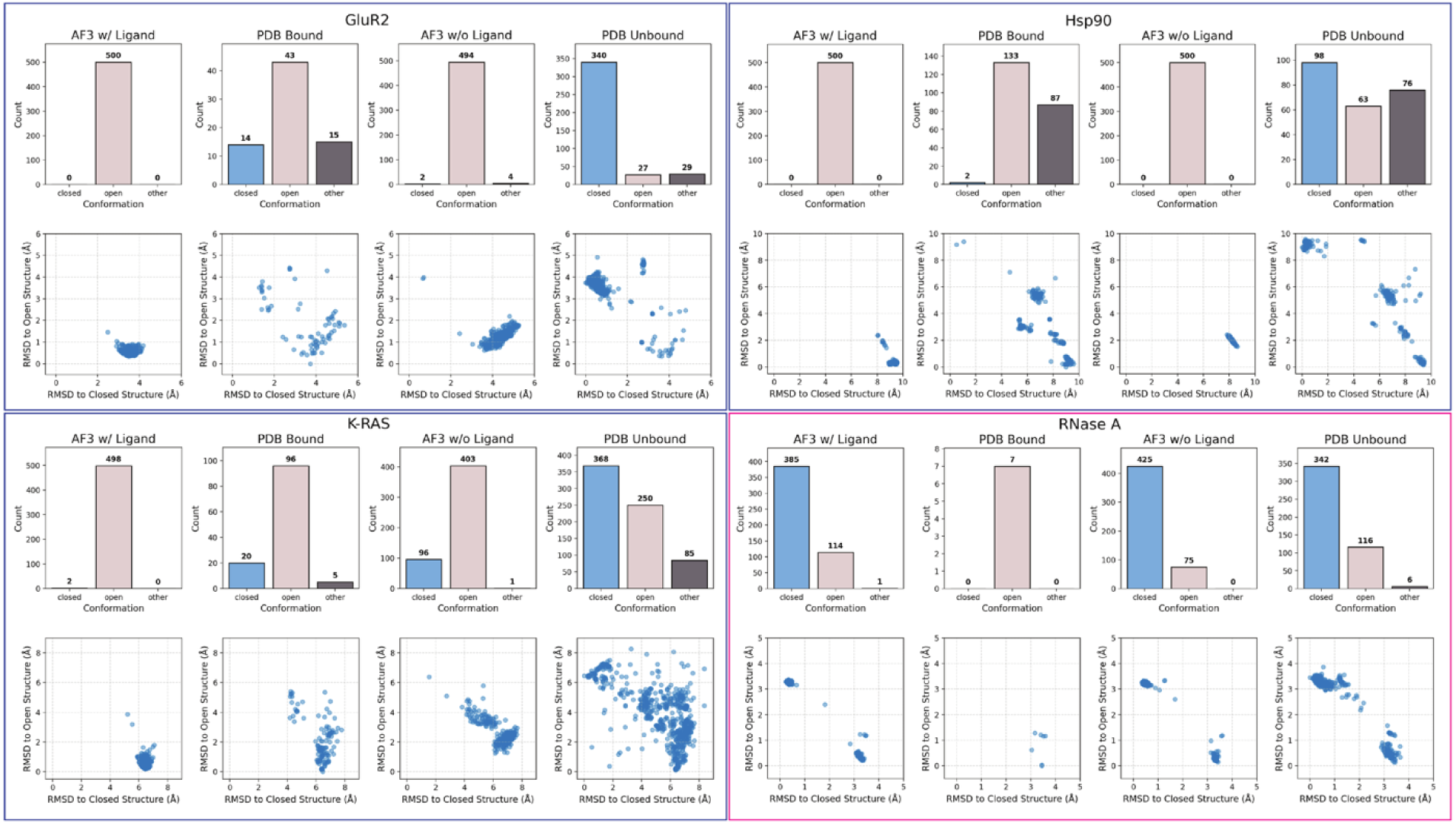
Comparison of bound and unbound structures with AF3 predictions with and without a ligand where the PDB distributions reveal an open and closed state, and AF3 accurately predicts one state. Group 1b (blue border) and Group 1c (magenta border) For Group 1b (blue border), AF3 with the ligand accurately predicts the bound state, while for Group 1c AF3 without the ligand accurately predicts the unbound state. Distributions of the structures (from left to right for each protein, AF3 with ligand, PDB bound, AF3 without ligand, PBD unbound) and RMSDs of the moving residues relative to the open and closed reference structures are shown.

For RNaseA, AF3 predicts the unbound distribution without the ligand correctly but not bound distribution with the ligand (Group 1c, Figure 2). The moving segment (His 119) is a single side chain in the middle of a beta-sheet. The bound PDB distribution is entirely open except that one of the structure has His 119 in both conformations, although there are very few bound structures. The unbound PDB distribution is mostly closed as expected although some structures are open and some have the His 119 in both conformations. AF3 with the ligand predicts mostly the closed conformation which may be an example of detrimental memorization.

For AR, and AmpC, the PDB distribution is close to correct (Group 2 proteins, Figure 3). The AF3 with the ligand incorrectly predicts the closed conformation, because AF3 places the ligand in the active site pocket (the orthosteric site) instead of the cryptic pocket. For AR, the bound PDB distribution is almost equally split between open and “other”, with “other” slightly outweighing open, but there are very few structures. Nonetheless, AF3 places the ligand at the dihydrotestosterone binding site, the active site, which is quite far from the cryptic site (Figure 4B). In fact, there are no models in which the ligand is placed in the cryptic pocket. Most of the “other” conformations in both the PDB distributions and the AF3 distributions are partially open with, for example, Met 734 open and the Lys 720 closed (e.g., 2PIO). In the reference bound structure, the cryptic pocket ligand (RB1) and the active site ligand (dihydrotestosterone (DHT)), are bound simultaneously. In fact, in all of the bound structures, DHT is occupying the active site while another ligand is in the cryptic pocket. It is possible that without the DHT present the RB1 would bind in the active site. In the closed reference structure, while the cryptic pocket is empty, a different ligand is bound in the active site. For AmpC, AF3 with and without the ligand predicts only the closed conformation. The ligand in the AF3 models is incorrectly placed in the nearby active site pocket, leaving the cryptic pocket empty and closed. In both these cases, this result could also be due to detrimental memorization since the number of open structures overall is much less than the number of closed structures.

**Figure 3.**
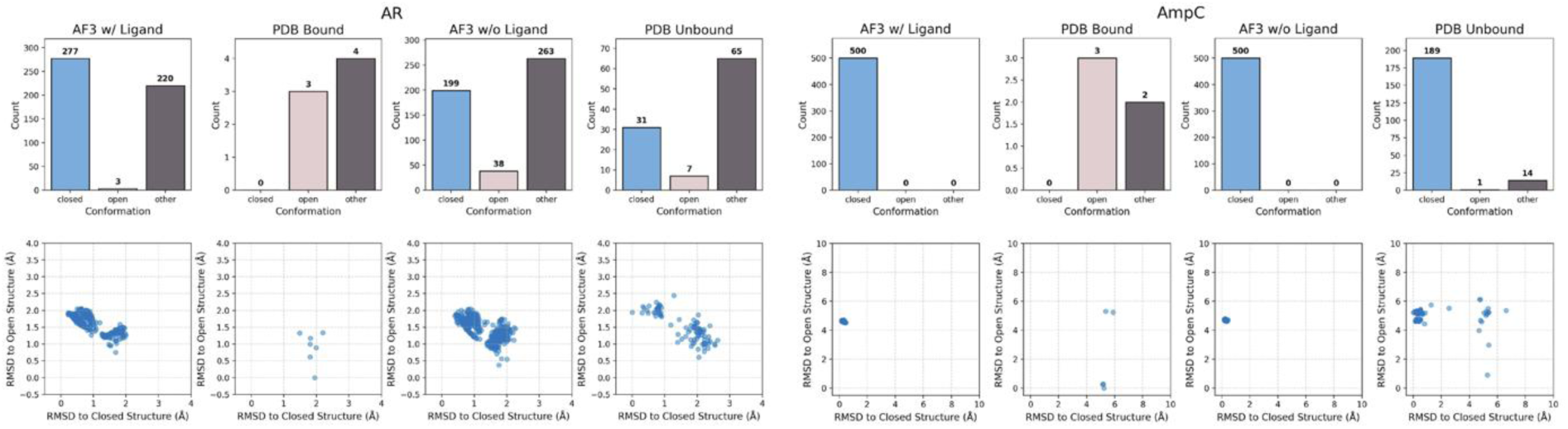
Comparison of bound and unbound structures with AF3 predictions with and without a ligand for Group 2 proteins, where the PDB distributions reveal multiple states including open and closed but AF3 place the ligand in the incorrect pocket. Distributions of the structures (from left to right for each protein, AF3 with ligand, PDB bound, AF3 without ligand, PBD unbound) and RMSDs of the moving residues relative to the open and closed reference structures are shown.

**Figure 4.**
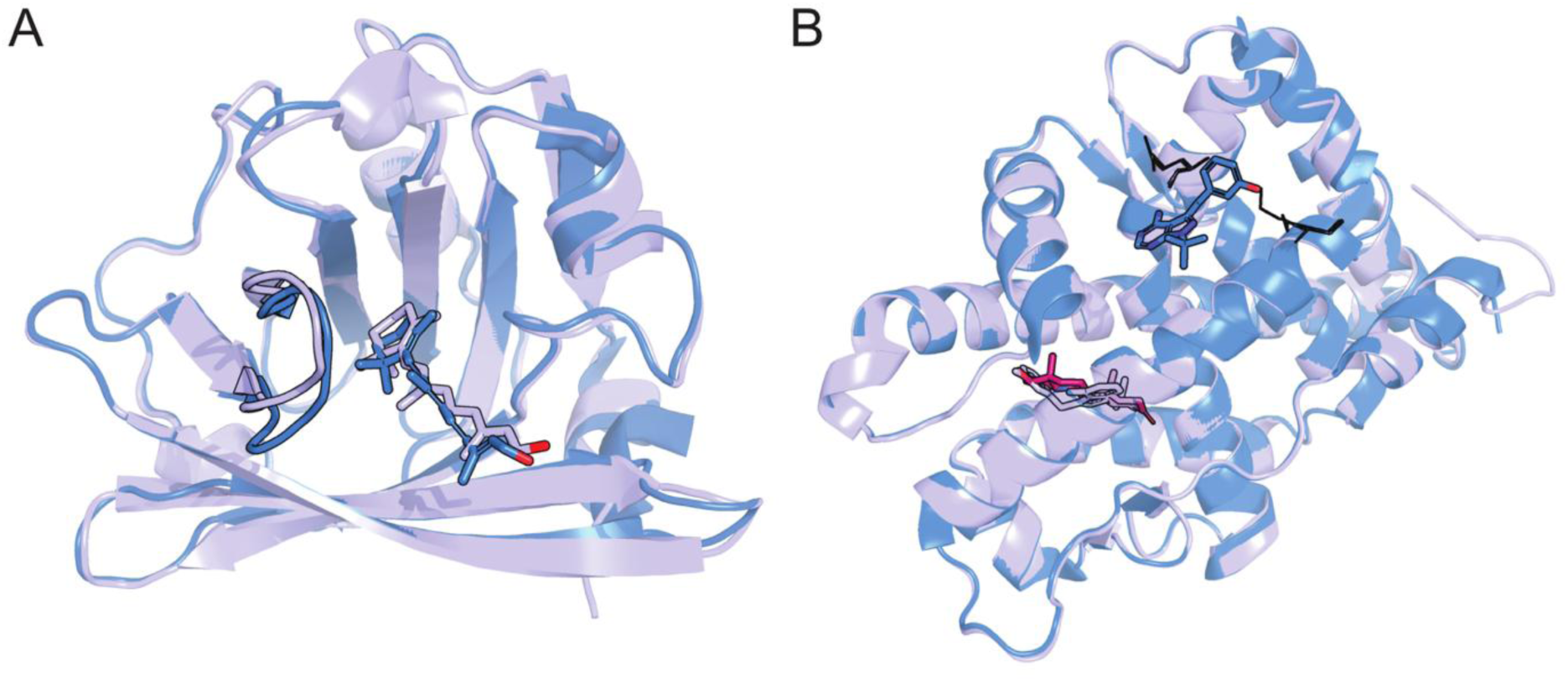
Superposition of the bound reference structure and the AF3-predicted structure co-folded with the reference ligand. Shown in (A) for BLG and in (B) for AR. The AF3 model with the ligand is shown colored by element with lilac carbons. The X-ray bound reference structure is shown colored by element with blue carbons. In (B) the DHT bound in the active site is shown in magenta.

For PKA, Thrombin, BACE (Group 3, Figure S2), there is almost a continuous distribution of loop conformations (the moving segment) from closed to open. For PKA, in the bound PDB distribution there is a wide range of open conformations, while for the unbound PDB distribution there is an almost continuous range of conformations spanning from closed to open. For Thrombin, there are distinct open and closed states, but a wide range of conformations around each. In the PDB unbound distribution, there are two clusters of closed conformations as well as many open conformations. Depending on the bound reference structure chosen, the conformations in the structures are either all classified as open (as in Figure S2) or all closed. For PKA and Thrombin, the cryptic pocket is open and accessible whether the ligand is bound or not. As a result, AF3 also predicts all open conformations. For BACE, most of the structures are classified as closed although there is a significant fraction of open conformations in both the bound and unbound PDB distributions. The AF3 predictions are all entirely closed which may reflect detrimental memorization. While there are a few instances of detrimental memorization, the effects over this test set seem to be less prevalent with AF3 than with AF2. ^24^

### Accuracy of ligand pose predicted by AF3

The reference ligand is predominantly positioned correctly by AF3 in 10/16 of the proteins (see Table 2, Figure 5A). For four other cases, the correct pocket is identified; for three of these (PKA, TEM, and K-Ras) some poses are also correct, for one (RNase A) only a very small number of correct poses are. Finally, for the last two proteins (AR and AmpC), the ligand, as discussed above, is positioned in the orthosteric site. There is a overall correlation between ligand predicted Local Distance Difference Test (pLDDT) score and the ligand pose correctness, since with the decrease of ligand pLDDT values of the ensemble the number of correct poses decreases as well (see Figure 5B).

**Figure 5.**
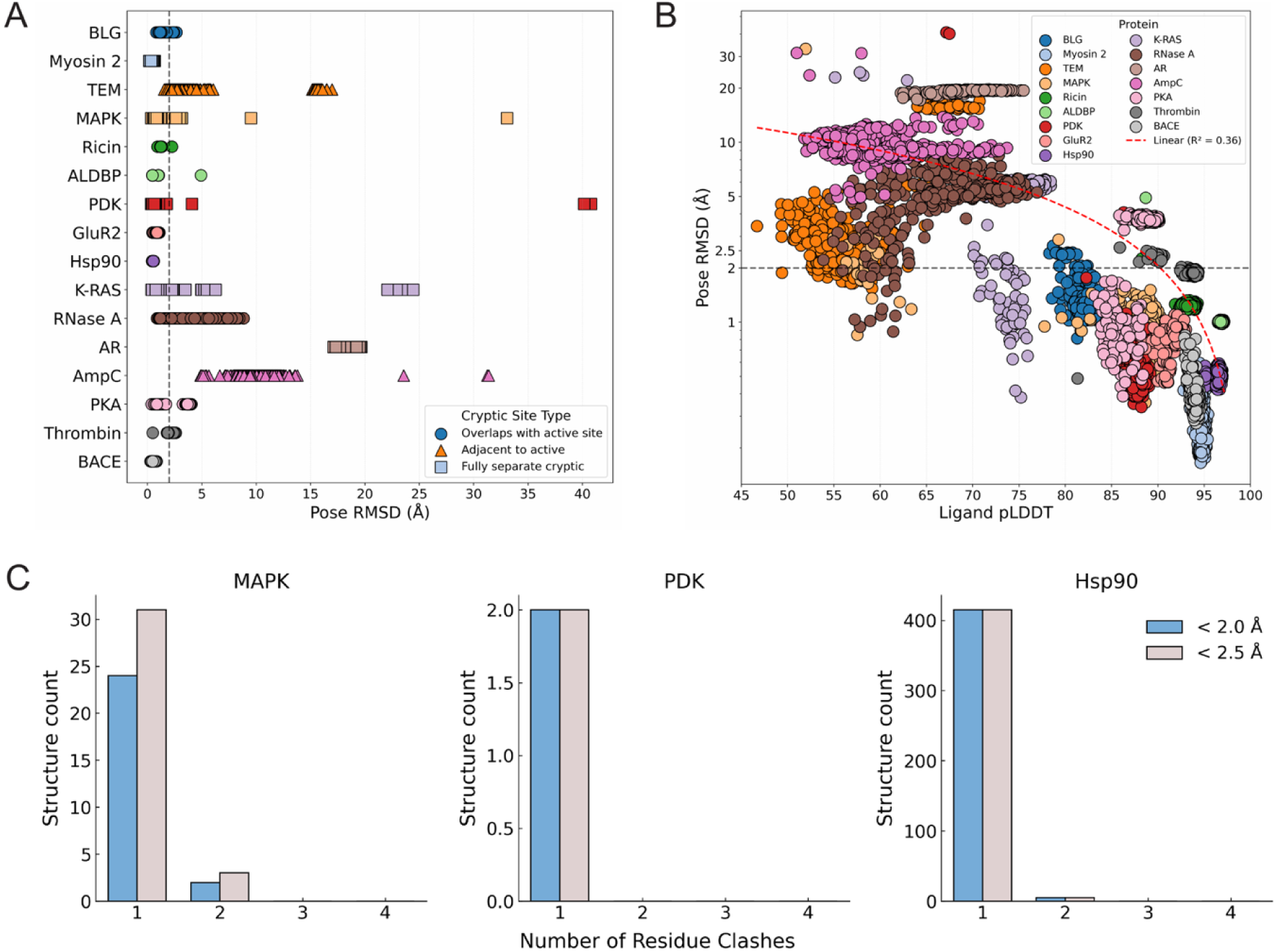
Evaluation of ligand pose accuracy in AF3 ensemble predictions. (A) Distribution of pose RMSD values across predicted ligand ensembles. (B) Relationship between pose RMSD and average ligand pLDDT confidence scores. (C) Frequency of residue–ligand steric clashes in AF3-predicted poses, shown only for cases where clashes were detected.

**Table 2.** Correct and near-correct ligand poses by protein for AF3 co-folded ensembles.

| Protein<sup>a</sup> | % $\leq 2$ Å | % $\leq 2.5$ Å | Correct Pocket |
| --- | --- | --- | --- |
| **BLG** | 94.2 | 99.8 | Yes |
| **Myosin 2** | 100 | 100 | Yes |
| **TEM** | 7.6 | 34.8 | Yes |
| **MAPK** | 95.8 | 98 | Yes |
| **Ricin** | 99.8 | 100 | Yes |
| **ALDBP** | 99.8 | 99.8 | Yes |
| **PDK** | 99.4 | 99.4 | Yes |
| **GluR2** | 100 | 100 | Yes |
| **Hsp90** | 100 | 100 | Yes |
| **K-Ras** | 9.6 | 11.4 | Yes |
| RNase A | 3.4 | 5.6 | Yes |
| AR | 0 | 0 | No |
| AmpC | 0 | 0 | No |
| **PKA** | 19.4 | 19.4 | Yes |
| **Thrombin** | 94.4 | 99.8 | Yes |
| **BACE** | 99.8 | 99.8 | Yes |

<sup>a</sup> In bold are the cases where AF3 with the ligand predicts the correct pocket conformation.

Notably, for three of the proteins, correct ligand poses are predicted in which the ligand still sterically clashes with the protein (see Figure 5C). For PDK this represents only two of the models predicted by AF3 (<1%). For MAPK it represents over 35 of the 500 models predicted by AF3 (still < 10%), and for Hsp90 it represents over 400 models predicted by AF3 (∼ 80%). In all three cases, the ligand pLDDT values were generally high; however, for MAPK and PDK, some predictions with lower pLDDT values corresponded to correct poses. Some but not all of the MAPK predictions with low ligand pLDDT scores had steric clashes, but all of the PDK models with clashes still have high ligand pLDDTs. Interestingly, ligand pLDDT for all Hsp90 models is high (> 95) despite most poses having a steric clash.

The data also indicate that if AF3 is run with a ligand, at least one conformation competent to bind a ligand in the cryptic site would be found for 14 out of 16 proteins in the test. This result is expected given that structures with ligands bound in the cryptic pocket exist and were in AF3 training set. It also suggests that, despite some inaccuracies, the ensemble of predicted structures should be useful for drug design purposes.

### The effect of the ligand on AF3 predictions

The choice of the ligand used for the AF3 prediction can sometimes affect the results, especially when the ligand binds to a conformation significantly populated in the PDB. For example, when the ligand-bound AF3 prediction for GluR2 is performed with the smaller ligand, glutamate, instead of the reference ligand, ATPO ^25^, the correct closed conformation is predicted. Glutamate is not large enough to open the cryptic pocket. Similarly, X-ray structures of TEM-Xe show Xe simultaneously bound in multiple sites including the cryptic pocket, all in the closed conformation. The AF3 predictions for TEM with one Xe bound also correctly predict the closed conformation but with the Xe in a different pocket. Interestingly, if we ask AF3 to predict TEM with 3Xe bound, it places all 3 Xe as overlapping in the cryptic pocket with the protein in the closed conformation. In our classification of bound vs. unbound structures in the PDB, the GluR2-glutamate and TEM-Xenon structures are all classified as unbound since there is no clash with the moving residues.

Furthermore, when AF3 is run for AR with both the cryptic site ligand (RB1) and the active site ligand (DHT) simultaneously, DHT is always predicted correctly to bind in the active site but the RB1 is still not placed correctly in the cryptic pocket. Instead, it is placed in a distinct third pocket on the surface of the protein (Figure 6). While the choice of the ligand has no effect on the MSA, since the model predicts the protein and ligand together, different input ligands will result in a different denoising path for the diffusion model.

**Figure 6.**
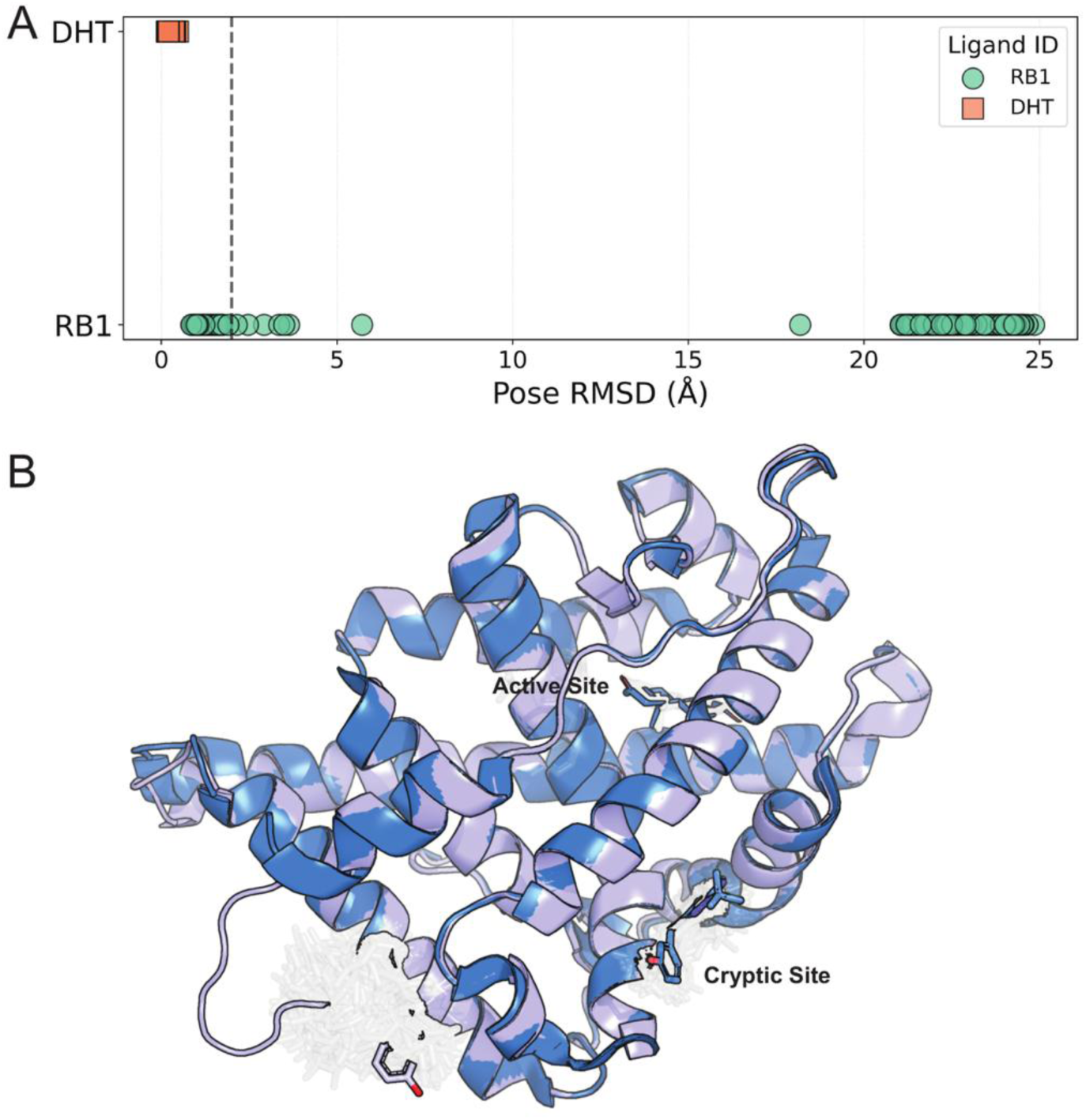
Evaluation of ligand pose accuracy in AF3 ensemble predictions for AR with the poses for DHT and RB1 predicted simultaneously. Shown is (A) is the distribution of pose RMSD values for DHT (active site ligand) and RB1 (cryptic site ligand) in the AF3 ensemble. In (B) are the AF3 models overlaid with reference bound structure (blue). The top ranked AF3 model by overall pLDDT (lilac) is shown in a ribbon representation; for the rest of the ensemble only the ligands are shown as sticks with transparency; the reference structure ligands are shown in blue sticks with DHT in top center and RB1 in lower right corner.

## Conclusion

AF3 is generally able to reproduce the scale of conformational change required for cryptic site formation upon ligand binding. That is, over our test set, when given a cryptic-site ligand for the protein, AF3 general predominantly predicts conformations competent to bind the ligand in the cryptic site. Conversely, when AF3 predicts the same protein structures without a ligand, the closed conformation generally dominates.

More often than not, AF3 tends to reproduce the dominant conformations observed in the PDB for the bound distributions with the ligand and for the PDB unbound distributions, even when there is a significant imbalance between the number of bound and unbound structures in the PBD (e.g., TEM, Ricin, ALDBP, GluR2, and K-Ras). Thus, while our results overall may reflect a bias toward memorized structural priors rather than flexible sampling of alternative conformations, the level of detrimental memorization appears to be reduced relative to what has previously been reported for AF2^24^.

The results clearly show that the same MSAs can be used to predict different conformations depending on the presence of a ligand. Somewhat surprisingly, the choice of the ligand given to AF3 can also significantly impact the predictions. We saw this with GluR2, for example, where when a ligand known to open the cryptic site was used the open conformation was predominantly predicted, while if glutamate (which binds at the same site but with the pocket closed) is used the site is correctly predicted as closed. We have also seen that AF3 has no ability to discriminate when two ligands are directly overlapping (as in the case of TEM with multiple Xe ions predicted to bind in essentially the exact same site); AF3 is unaware that that is physically impossible. In the case of AR, when AF3 is given the active site ligand and the cryptic site ligand simultaneously, it rarely places both ligands in the correct pocket.

For all but two of the proteins, in our test set we should that AF3 was able to produce at least a few models with the ligand correctly position; the exceptions were AR and AmpC where AF3 incorrectly place the cryptic site ligand in the active site. Furthermore, we have also shown that ligand pLDDT scores over the AF3 structural models in the ensemble correlate somewhat with the ligand pose RMSD (with R^2^ = 0.35). That is, models with lower ligand RMSDs tend to have higher ligand pLDDTs. For three of the proteins (MAPK, PDK, and Hsp90), even when the ligand was placed correctly there could be clashes with the protein; these represented a minor number of models for MAPK and PDK but ∼80% for Hsp90. The ligand pLDDTs for Hsp90 are high (ranging from about 95 to 97). Again, this reflects the fact the AF3 is not fully aware of the physical constraints. The variability in ligand position underscores the importance of generating ensembles of ligand-cofolded predictions using multiple seeds to maximize the likelihood of obtaining a correct binding mode. Despite the limitations, using AF3 with extensive sampling to generate structural ensembles can reveal cryptic pockets that may be useful for drug discovery. The challenge will be in identifying those conformations with actionable cryptic pockets that are revealed among the many conformations generated.

## Methods

### Test set description

The test set utilized for this study was the same as in Lazou et al.^24^ All proteins in the test set (see Table 1) originate from the CryptoSite database^4^, except for K-Ras. In assembling CryptoSite, Sali and colleagues compared ligand-bound structures with the corresponding unbound structures from the PDB. They applied pocket-detection algorithms and retained only those pairs in which the pocket was minimal or absent in the unbound state but underwent a substantial conformational change - either expansion or closure - to accommodate the ligand upon binding. For the 15 CryptoSite proteins, these criteria defined the unbound and bound reference conformations and the moving segment. Typically, the ligand-bound structure also represents the open state, as binding exposes or enlarges the pocket. For two proteins—thrombin and β -secretase—the binding region is too open or shallow in the unbound conformation, and ligand binding instead induces a closing of the site. In these cases, the bound state corresponds to the closed rather than the open reference. For K-Ras, the pocket and the relevant PDB entries were taken from a study by Fesik and co-workers^26^, which showed that the site becomes accessible when a small molecule binds. Overall, the bound and unbound structures provide complementary reference states, with the conformational transitions detailed in Table 1. Although these changes may involve multiple loops or even entire domains, in each case a single loop movement or a side-chain rearrangement was sufficient to capture the main structural difference between the two states.

### Protein Data Bank distributions

For each of the 16 cases, we assembled a set of experimental structures and categorized them into bound and unbound states. Experimental structures were retrieved from the PDB using the RCSB API by searching for entries with ≥90% sequence identity to the AF reference PDB ID (Table 1). ^27,28^ Binding was determined through geometric criteria. A structure was considered bound if the distance between the ligand’s geometric center in the reference structure (ref_center) and the corresponding center in the test structure (test_center) was less than 5.0 Å, indicating that the ligand occupied a similar spatial position. Alternatively, binding was also assigned if any atom in the reference ligand was within 2.0 Å of any atom in the test ligand, a stricter overlap threshold designed to capture close atomic contacts. In cases where the cryptic site was adjacent to the active site, an additional criterion was applied: a structure was considered bound only if the ligand clashed with residues in the moving segment of the unbound structure. Clashes were calculated using a KDTree-based algorithm: for each atom, potential interacting atoms were identified and classified as clashing if their interatomic distance to the original atom was less than 0.63 of the sum of their van der Waals radii.

### AF3 predictions

AF3 code was obtained from the official AlphaFold3 repository (https://github.com/google-deepmind/alphafold3), and model weights were downloaded from Google DeepMind with their permission. Structure predictions were performed by first running the MSA search with MMseqs^29,30^, followed by converting the resulting alignments into JSON format using the alphafold3_tools repository (https://github.com/cddlab/alphafold3_tools). No structural templates were used, and predictions were generated with 100 random seeds, which resulted in total 500 models. For each of the 16 cases, the predictions were performed with and without a ligand present. For the ligand-bound predictions, the SMILES corresponding to the ligand listed for each case in Table 1 was used unless otherwise stated.

### Analysis of conformations

For each of the 16 cases, the two sets of structures (with and without ligand) were aligned to both the reference closed and open states. The RMSD of the moving residues, as defined in Table 1, was then calculated. For each structure in the PDB distribution and for each model in the AF3 ensemble, the RMSD of the moving segment from both the open and the closed reference structures were determined. The moving segment was excluded during the alignment to the reference structures and the RMSD was only calculated for the moving residues. For proteins with moving loop segments, only alpha carbons are considered in the RMSD calculations, while for systems with individual side chains identified as moving segments, all-atom RMSD is calculated. In Figues 1–3 and S1, the structures are represented in a 2D local coordinate system that shows the RMSD of the moving segment from the closed reference structure on the X axis and the RMSD of the moving segment from the open reference structure on the Y axis. To classify each structure as open, closed, or “other”, we first computed the RMSD between the open and closed reference structures and used half of this value as the cutoff. Structures with an RMSD to the open reference below this threshold were assigned to the open state, while those within the threshold of the closed reference were assigned to the closed state. Structures that did not meet either condition were classified as “other” state. For regions defined by continuous amino acid segments, we calculated RMSD using Cα atoms, whereas for regions defined by side chains, all heavy atoms were used. In addition, for cases where the classification of the PDB structures did not result in open conformations dominant for bound and closed dominant for unbound as expected (K-Ras and Ricin), we added the criterion that any structure or model for which that residue (the moving segment) would clash with the reference ligand (in the aligned reference structure) would be classified as closed.

## Acknowledgements

This work was supported in part by grant R35GM118078 from the National Institute of General Medical Sciences.

## Author contributions

M.L. and F.T. conducted the research. M.L. and D.J.-M. wrote the manuscript. S.V. supervised the research. D.J.-M. conceptualized the project and supervised the research.

## Competing interests

The authors declare no competing interest.

## Data and code availability

All structures utilized in this work are summarized in Table 1. Model weights for AF3 were downloaded from Google DeepMind. Scripts for analysis are available on GitHub at https://github.com/J-MLab/af3_cryptic.

## Abbreviations

BLG: Beta-lactoglobulin
TEM: TEM β-lactamase
MAPK: Mitogen-Activated Protein Kinase
ALDBP: Adipocyte Lipid Droplet Binding Protein
PDK: Pyruvate Dehydrogenase Kinase
GluR2: Glutamate receptor 2
Hsp90: Heat Shock Protein 90
RNase A: Ribonuclease A
AR: Androgen receptor
AmpC: AmpC b-lactamase
PKA: cAMP-dependent protein kinase
BACE: β-Secretase

## Notes

### Competing Interest Statement

The authors have declared no competing interest.

