## Supplemental Figures S1 and S2. for "The Influence of Ligands on AlphaFold3 Prediction of Cryptic Pockets"

This file includes:

**Figure S1**. Superposition of the bound reference structure and the top ranked AF3-predicted structure co-folded with the reference ligand.

**Figure S2**. Comparison of bound and unbound structures with AF3 predictions made with and without a ligand for Group 3.

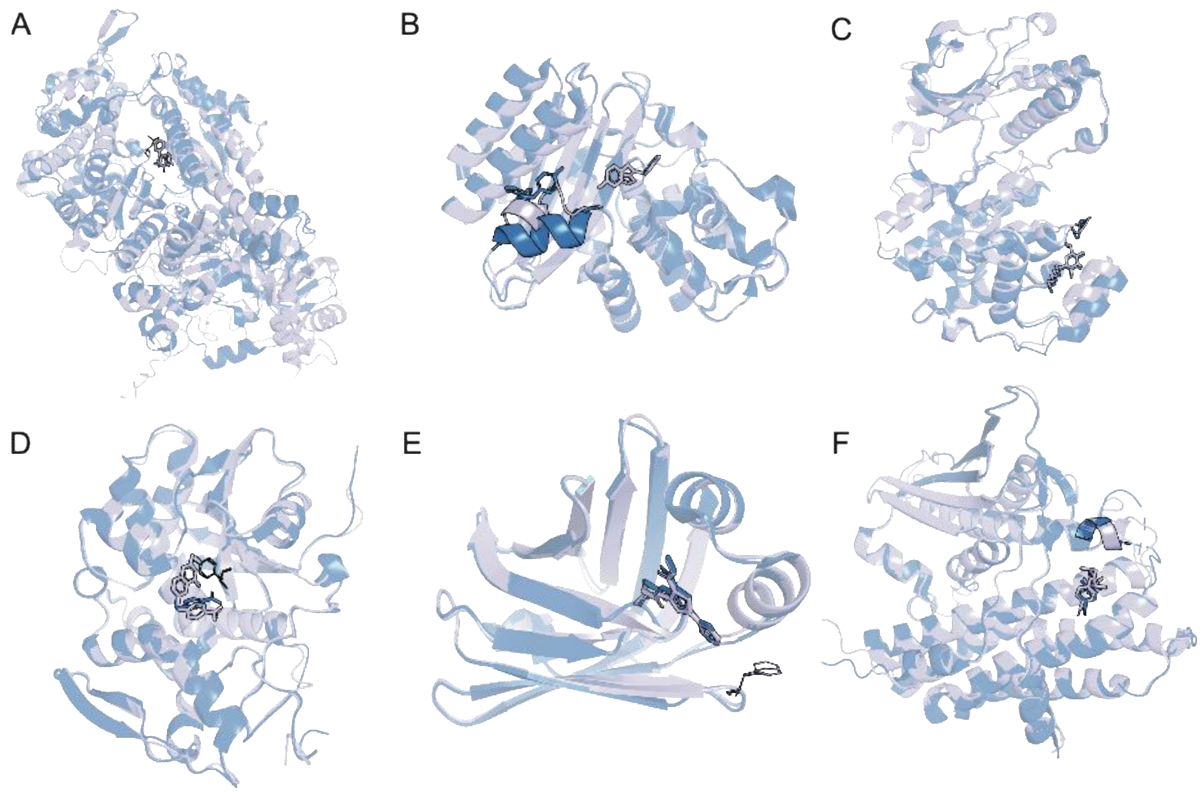


**Figure S1. Superposition of the bound reference structure and the top ranked AF3-predicted structure co-folded with the reference ligand.** Shown in (A) Myosin II, in (B) TEM, in (C) MAPK, in (D) Ricin, in (E) ALDBP, and in (F) PDB. The AF3 model with the ligand is shown colored by element with lilac carbons. The X-ray bound reference structure is shown colored by element with blue carbons. Moving segment is shown with lines or no transparency setting.

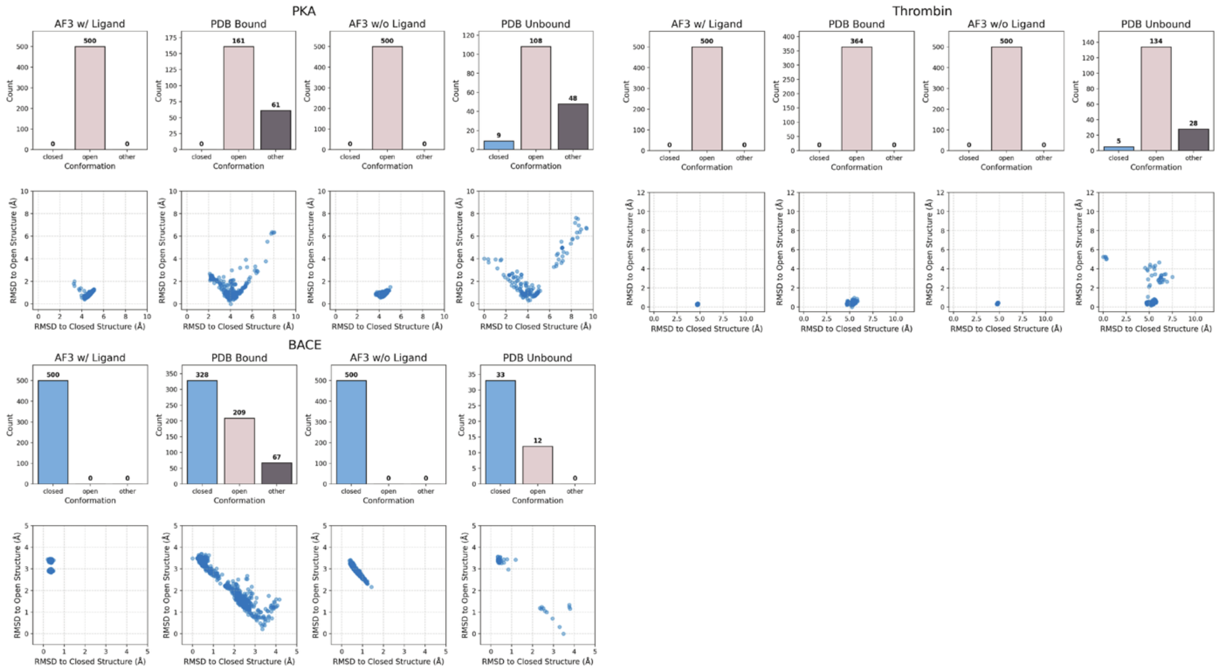


**Figure S2. Comparison of bound and unbound structures with AF3 predictions with and without a ligand for Group 3 proteins, where the PDB distributions reveal one predominant state due to the structures spanning the full range from open to closed, and AF3 predicts the dominant state.** Distributions of the structures (from left to right for each protein, AF3 with ligand, PDB bound, AF3 without ligand, PBD unbound) and RMSDs of the moving residues relative to the open and closed reference structures are shown.
